# Does voluntary wheel running exist in Neotropical wild mammals?

**DOI:** 10.1101/409409

**Authors:** Peter van Lunteren, Marnix A. Groenewold, Gabor Pozsgai, Joseph Sarvary

**Affiliations:** These authors should be considered joint first author, as they contributed equally to this work; Fundacion Para La Tierra, Mariscal Estigarribia 321 c/Teniente Capurro, Pilar, Ñeembucú, Paraguay; State Key Laboratory of Ecological Pest Control for Fujian and Taiwan Crops, Fujian Agriculture and Forestry University, Fuzhou 350002, China

**Keywords:** Neotropics, stereotypic behavior, synurbization, urbanization, wheel running, wild environment

## Abstract

Running wheels are frequently used to improve the welfare of captive animals, increase environmental enrichment, and, by doing so, reduce stereotypic behaviors. It is, however, still debated whether or not wheel running itself is a stereotypy. New evidence emerged when Meijer and Robbers (2014, Proc. Royal Soc. B) reported voluntary wheel running of wild animals in the Netherlands. Since stereotypic behaviors are exclusively attributed to captive animals, the occurrence of wheel running in the wild suggests that this behavior is non-stereotypic. Our study explores that same line of investigation, examining whether wild animals will voluntarily use running wheels in a natural area in Paraguay in comparison to the urban and semi-urban settings in the Netherlands. Of the 1857 small mammal visits we recorded, only two occasions showed evidence of what could be considered as wheel running behavior; over hundredfold fewer than previously reported. The potential reasons for the observed difference in wheel running activity, such as different species pool or seasonality, are discussed. The difference, however, is likely to be due to the much lower probability of Neotropical mammals in a remote natural site encountering man-made objects and experiencing urbanization-related behavioral patterns. Additionally, in the light of our findings, we review the definition of wheel running as a stereotypic behavior.

## Introduction

Across the world, animal welfare has become a topic of increasing ethical and political concern (Mason and Mendl, 1993; Sneddon et al., 2016). The welfare of captive animals depends on many factors, of which one of the most important is the stimuli in the environment in which animals live (Clark et al., 1997). Unsuitable, stimulus-poor environments can lead to chronic stress in captive animals (Terio et al., 2004; Morgan and Tromborg, 2007), which in turn can cause the development of stereotypic behaviors (Würbel, 2001). Stereotypies are defined as abnormal, repetitive, invariant, and apparently functionless behaviors (Mason, 1991) and can result in self-injury or reduced fitness (Garner and Mason, 2002). Wheel running behavior is thought to represent exploratory migration, and, in captivity, attempts to escape (Mather, 1981). Another explanation for the motivational basis of wheel running could be thermoregulatory needs (Janik and Mrosovsky, 1993). Wheel running is considered self-reinforcing and perceived by captive animals as ‘important’ (Sherwin, 1998). It additionally has documented beneficial effects on captive animals’ development (Ehninger and Kempermann, 2003; Rhodes et al., 2003) and physiological systems (Colbert et al., 2006; Werner et al., 2009). Wheel running increases when environmental quality declines (Ödberg, 1987; Powell et al., 1999; Shyne, 2006) and is inversely correlated with stereotypic behaviors, including the stereotypy known as ‘bar mouthing’ (Hansen and Berthelsen, 2000; Gebhardt-Henrich et al., 2005; Richter et al., 2008). Consequently, running wheels are often used to increase environmental richness (Olsson and Dahlborn, 2002), thus improving animal welfare, and decreasing stereotypic behavior for captive animals (Würbel et al., 1998; Hansen and Berthelsen, 2000).

However, rodents with excessive wheel running behavior often show brain malfunction (Mathes et al., 2010) and it is still much debated as to whether or not wheel running itself is a stereotypic behavior (Mason, 1991; Sherwin, 1998; Mason and Latham, 2004; Richter et al., 2014). Since wild, free-living animals are not thought to show stereotypic behaviors, the occurrence of wheel running in wild environments could unravel this question and improve our understanding on its significance for behavioral science.

To the extent of our knowledge, wheel running behavior in wild environments has only been tested by Meijer and Robbers (2014). These authors demonstrated that wheel running was voluntary for wild animals in the Netherlands and argued that “wheel running does not fit well within the definition of a stereotypy”. Their study sites — an urban and a dune habitat, however, were more characterized by urban, rather than wild areas, where animals have a greater chance of encountering humans or man-made objects. Therefore, Meijer and Robbers (2014) do not provide convincing evidence that truly wild animals use running wheels voluntarily. Thus, we believe that wheel running behavior in non-urban habitats should be further investigated and the knowledge gaps in this poorly explored topic filled.

This experiment rethinks the work carried out by Meijer and Robbers (2014) with the aim of investigating whether wild mammals found in a natural site in eastern Paraguay show voluntary wheel running behavior similar to that reported in the Netherlands. In the light of new evidence, we also contribute to the debate whether or not wheel running fits the definition of stereotypic behavior.

## Materials and Methods

The study was conducted at Para La Tierra Ecological Station, located at Rancho Laguna Blanca in Paraguay, South America (S 23°48’45.4”, W 56°17’41.7”). This experiment was based on the methodology described by Meijer and Robbers (2014), with some adjustments made to adapt to local conditions. Running wheels were set up at one location in a secondary dry forest, two locations in a South American tropical savannah (cerrado, figure 1*a*), and five locations in a semi-deciduous transitional humid-dry gallery forest (transitional forest, figure 1*b*) which is based around a 157 ha freshwater lake (Smith et al., 2016). The nearest human village, Santa Barbara, with 2500 inhabitants (Joseph Sarvary, personal communication, Oct, 2016), was approximately 3.8 km from the closest running wheel. The eight individual locations were established at least 445 m from any other study location. Since the typical home ranges of small mammals of interest occurring at Laguna Blanca (i.e., rats, mice, and opossums) do not exceed 1 ha (Mikesic and Drickamer, 1992; de Almeida et al., 2008; Umetsu et al., 2008), the minimum distance between our sampling locations provided ample area to minimize the chance of one individual visiting two or more wheels (a mean of 33.6 ± 26.3 ha (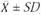) non-overlapping area, with a smallest being 17.3 ha, figure 2). Various running wheel constructions, wheel diameters, food preferences, and types of protective cages were tested in a six-week period of preliminary study (data not presented). In the final design, all running wheels were 30.5 cm in diameter and were built on-site using locally available plastic bowls and bearings (figure 3). The wheels were designed to be light enough to enable small animals to move them and visiting animals were able to enter and leave the site without any restriction. The use of a protective cage, as used by Meijer and Robbers (2014), was found to be unnecessary, since food was generally consumed by animals of interest. Moreover, no signs of predator attacks were noted and no substantial differences in visits or wheel movements were observed between cage and no cage controls in the preliminary study. Preliminary experiments also showed the presence of a relatively larger mammal species, the white-eared opossum *Didelphis albiventris* (Lund, 1840), with an average body length of approximately 34 cm, excluding the tail (Smith, 2007). Since opossums are known to run in wheels in captivity (Cummins, 2006), we adjusted the wheel size to enable these, as well as other small mammals, to enter and move the wheel with ease. A mixture of oats and local fruit, placed approximately 20 cm in front of the wheel, was used to attract animals. However, in order to test the influence of food availability, we stopped providing food at all sites simultaneously after six weeks (equaling to 48 wheel-weeks with bait). The unbaited period lasted an additional five weeks (equaling to 40 wheel-weeks), which resulted in an *overall observation period* of 88 wheel-weeks. Due to adverse weather events, camera malfunctions, or other circumstances that made data collection impossible, a number of days were removed from this *overall observation period*, resulting in a *net observation period* of 50 wheel-weeks. Data were obtained from early April 2016 to mid-May 2016 and from mid-May 2016 to late June 2016 for the baited and unbaited period, respectively, by using Bushnell 8MP Trophy Cam and Crenova RD1000 trail cameras. Similarly to those used by Meijer and Robbers (2014), these cameras operate with infrared motion detection that does not interfere with the mammals’ behavior (Jacobs et al., 1991).

**Figure 1.**
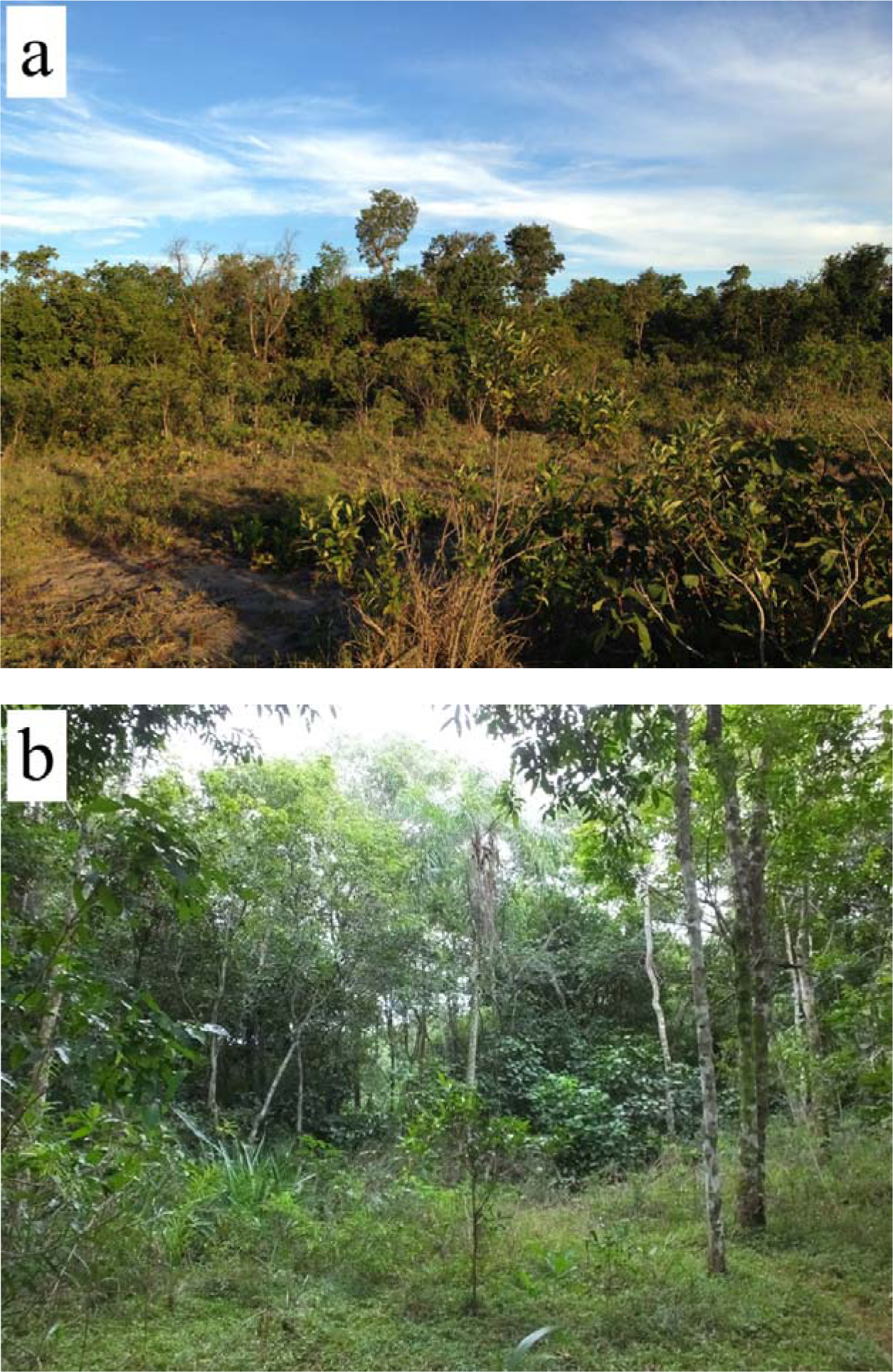
Photographs of the cerrado (a) and transitional forest (b), two habitat types of Rancho Laguna Blanca, Paraguay, where the wheels were placed. Photographed June, 2016.

**Figure 2.**
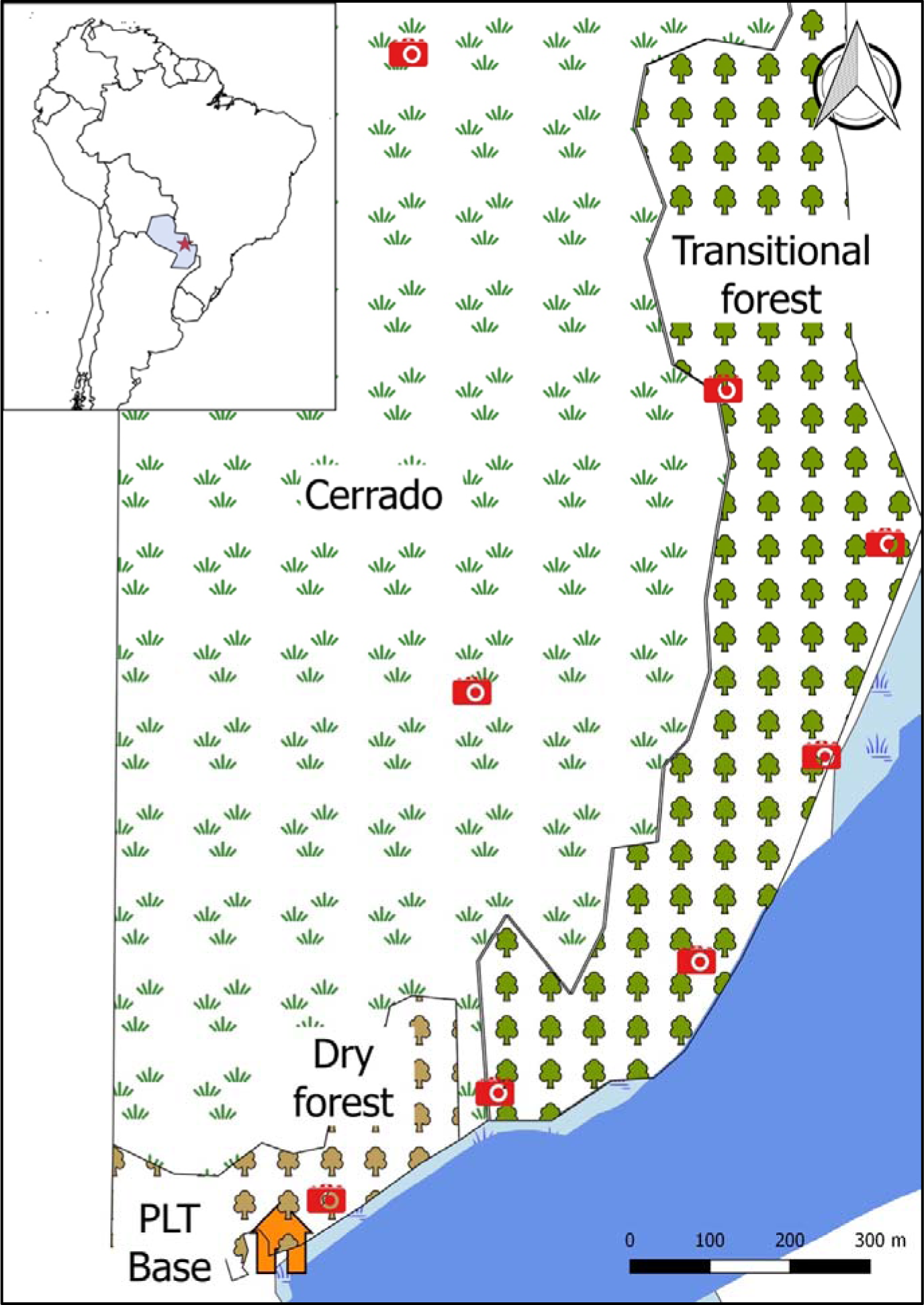
Map of the location where the wheels were placed. The three habitats (dry forest, transitional forest, and cerrado) are marked with brown tree, green tree and grass turf symbols, respectively. Camera icons indicate the locations of the traps, the orange house icon indicates Fundacion Para La Tierra Estacion Ecologica, and dark and light blue colors indicate lake and the wet lakeshore area, respectively. Rancho Laguna Blanca is marked with a red star on the overview map. Created June, 2016.

**Figure 3.**
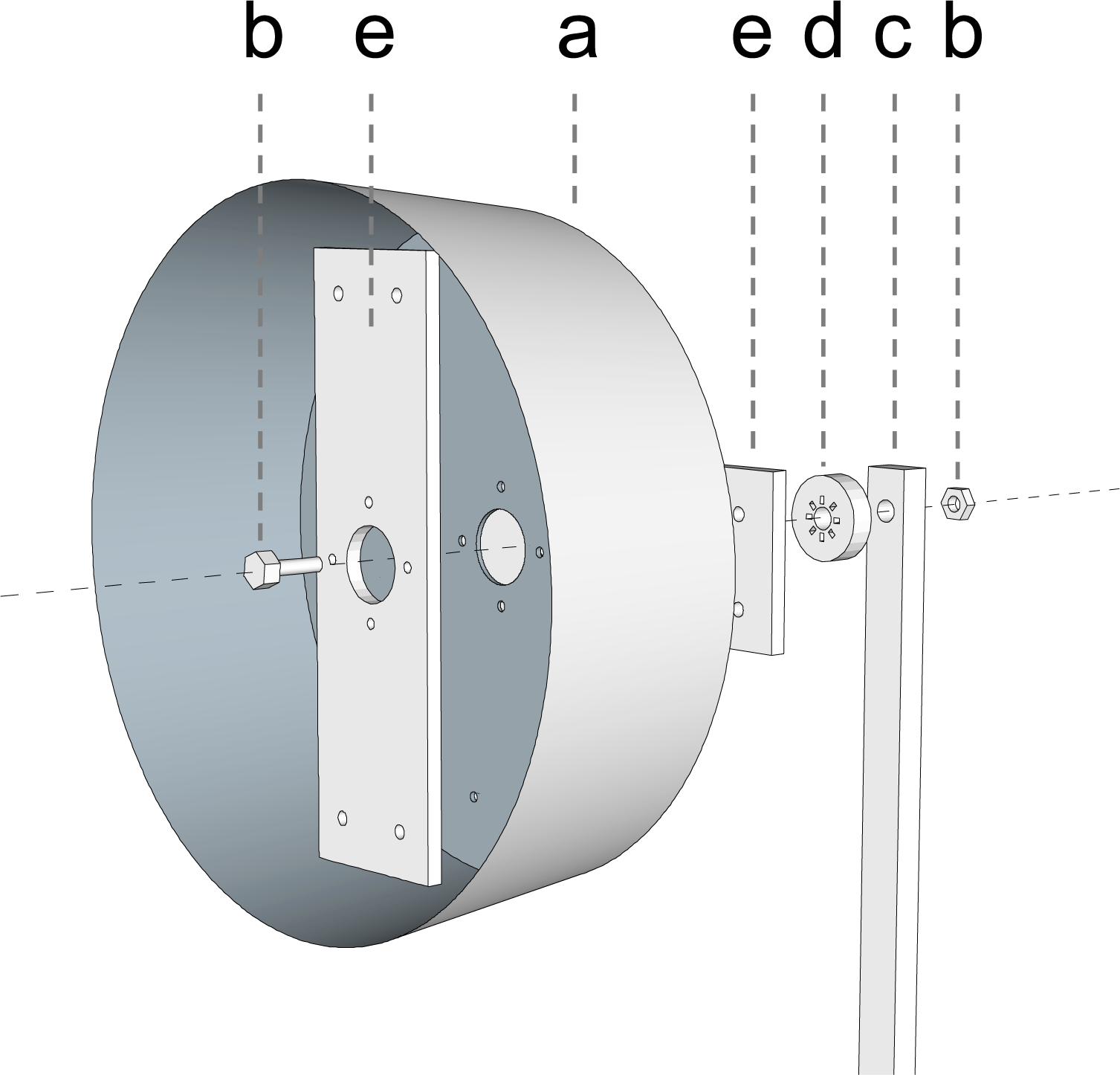
The schematic design of a running wheel used in the experiment. A plastic wheel (a) was attached with a bolt and a nut (b) to a wooden beam (c), between which bearings (d) were placed to minimize the friction. To maximize the stability, two small wooden planks (e) were attached crosswise to the front and back side of the wheel. Sketch constructed June, 2016.

Videos were examined for the presence of three types of wheel movement: 1) wheel running, defined as non-haphazardous directional running inside the wheel causing at least half a rotation; 2) wheel movement from inside the wheel (WMI), described as any wheel movement caused by an animal using all four legs inside the wheel that was not wheel running, and 3) wheel movement from outside the wheel (WMO), wheel movements caused by animals with fewer than all legs within the wheel (for examples see Supplementary Data S2 – S5). The duration of activity was measured in seconds, while the number of spins are estimated to the nearest half rotation, with a minimum initial value of half a rotation (equaling a distance covered of 48 cm). Only wheel movement caused by small mammals (e.g., rats, mice, and opossums) were included in the analysis. Other animal visits (e.g., armadillos, birds, insects etc.) were rare, and none of these animals were observed running in the wheel. However, the number of visiting animals included in this study — despite the exclusion — is abundant and sufficient for comparison.

## Results

During the *net observation period* 1857 animal visits (1591 rats; 149 mice; 117 opossums) were recorded, of which 1775 (1532 rats; 135 mice; 108 opossums) in the baited and 82 (59 rats; 14 mice; 9 opossums) within the unbaited period. The number of animal visits declined substantially after we stopped providing food, from an average of 46.0 visits/week/wheel during the baited period to 7.7 visits/week/wheel during the unbaited period. WMI was recorded 59 times (51 times in the baited and 8 in the unbaited period), while WMO was recorded 101 times (93 and 8 times during the baited and unbaited period, respectively). Subsequent relative values are 0.029 WMI/visit and 0.052 WMO/visit for the baited period, whilst the unbaited period resulted in the values of 0.098 WMI/visit and 0.098 WMO/visit.

Even though our preliminary data are not presented in this paper, during this period an unusual behavior was observed which is worth reporting here. A steering-like movement from outside of the wheel was observed 28 times with an average duration of 4.7 ± 4.0 s (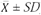) by opossums (Didelphidae, suspected *Marmosa* spp.), which, to our knowledge, has never been documented before (see Supplementary Data S6). We call this new behavior ‘wheel pulling’ and defined it as a wheel movement caused by an animal actively pulling the rim, while being outside the wheel. Since the *Marmosa* genus consists of arboreal animals, investigating whether ‘wheel pulling’ by climbers could be a substitute for wheel running is a promising area for future research. If this hypothesis proves to be correct, the captive conditions of climber species kept in limited spaces might also be substantially improved by means of re-designed running wheels, since the presence of these wheels could have the same beneficial effects for the welfare of climber species.

Wheel running behavior in Paraguay was only observed twice. One occurred in the transitional forest during the baited period and involved an unidentified opossum (Didelphidae, suspected grey short-tailed opossum *Monodelphis domesitca* (Wagner, 1842)) that was running for 9 s, covering a distance of 1.9 m. The second wheel running case was also recorded in the transitional forest but during the unbaited period, whereas an unidentified rat ran for 9 s, covering a distance of 1 m (see Supplementary Data S1 – S3).

## Discussion

In this study, following the research of Meijer and Robbers (2014), we investigated whether wild animals in Paraguay voluntarily use running wheels, a piece of equipment which is normally associated with mammals held in captivity. In order to allow comparison between Paraguay and the Netherlands we calculated runs per wheel per month for both studies. Meijer and Robbers (2014) recorded 44 months (equal to 191 wheel-weeks) in their baited and 17 months (equal to 74 wheel-weeks) in their unbaited period. Consequently, by dividing the total amount of wheel running cases by the *net observation period*, Meijer and Robbers (2014) observed wheel running 127 times more frequently in the Netherlands than we did in Paraguay with 22.0 and 0.2 wheel running cases per month per wheel, respectively. When comparing solely the baited phases, wheel running was observed 259 times more frequently in the Netherlands, with 28.8 wheel running cases per month per wheel versus the 0.1 wheel running case per month per wheel in Paraguay.

Additionally, we recorded a substantial difference between the two studies for the distance covered and the speed in which small mammals ran. In the Netherlands, mice ran for more than a minute in 20% of the cases, for a maximum of 18 minutes and with a maximum speed of 5.7 km/h (Meijer and Robbers, 2014), whilst in our observations wheel running did not exceed 9 s or a velocity of 0.8 km/h. The mice described by Meijer and Robbers (2014) “only ran in the wheels and never walked slowly”, as they did in our wheels. Therefore, not only the frequency with which wheel running occurs, but also the duration, velocity, and distance covered during wheel running cases differ considerably. Considering the short duration and the rare frequency of our recordings, it is debatable whether the two incidents we classified as wheel running truly show deliberate wheel running actions or might just be artefacts of mammals attempting to navigate their environment.

### Potential causes of the observed difference

We identified four variables that may have potential to influence wheel running activity; 1) the experimental set-up, 2) climate and seasonality, 3) the small mammal species pool, and 4) the probability with which animals are likely to encounter human objects in the wild.

#### Experimental set-up

Since the differences in wheel running activity, length, and velocity between the Netherlands and Paraguay remained substantial after standardizing for observation time (see Materials and Methods), it is unlikely that all these differences were caused solely by the difference in the length of the observation period. Moreover, since videos of all sites, including the preliminary phase, demonstrate animal interaction with the wheels from the first week, it is realistic to assume that the length of habituation period to the novel object was negligible, and the difference in the length of the study period did not substantially skew the results. Other main differences in our experimental set-up compared with that of Meijer and Robbers (2014) were the lack of protective cages, the slightly bigger wheel sizes (i.e., diameter of 30.5 cm vs. 24 cm), and the shorter study period in Paraguay. Based on preliminary tests, the use of cages was unnecessary since no substantial difference in visits was found and no behavioral variation was observed between animals visiting caged wheels and those visiting the controls. A slightly larger running wheel was chosen to adapt to local fauna but the weight of the wheels was not exclusionary to even the lightest bodied mammals observed visiting the wheel sites. Some variation in the observed wheel running activity could be explained by the differences between our definition of wheel running and the one used by Meijer and Robbers (2014) with the latter including ‘wheel running’ slugs and frogs in their analysis. This might imply a less strict definition used in the Netherlands. Meijer and Robbers (2014), unfortunately, did not provide their definition for wheel running. Our data, however, differentiate between animals running in the wheel and those moving the wheel from inside, while not properly running (i.e., 59 times, e.g., see Supplementary Data S4). This difference in definitions may cause some inconsistency, thus it may be desirable to re-categorize behaviors with standardized definitions. This dissimilarity, however, is unlikely to cause any significant difference since the wheel running cases in the Netherlands differed considerably from the ones in Paraguay in terms of duration, velocity, and distance covered. In summary, the methodological differences are sufficiently minor, thus they are unlikely to explain the observed low frequency of wheel running activity in Paraguay. From the perspective of experimental set-up, our results are suitable for comparison with Meijer and Robbers’ (2014) research.

#### Seasonality

Similarly to the length of the study, timing might also influence the results. Seasonality affects rodent behavior (O’Farrel, 1974) and has also been documented to influence wheel running activity (Wollnik et al., 1991). Wheel runs were recorded year-round in the Netherlands, with the frequency of runs increasing in late spring, with a peak in summer and autumn. Our study ran from April to June, which are the first few months of the dry season in Paraguay and, in temperature and precipitation, similar to mid-to-late summer in the Netherlands (The World Bank Group, 2017a). Our study period, therefore, corresponds with the peak of wheel running activity in the Netherlands. Furthermore, although very little is known about their phenology, Neotropical small mammals are active throughout the year (Fleming, 1971). In the present study, the number of visits at our wheels prove that there was ample animal activity during the recording period to have provided opportunity to observe wheel running were it to occur naturally. For these reasons, we suggest that seasonal differences are unlikely to have caused the observed disparity in wheel running activity between the two studies.

#### Species pool

Due to the substantial difference in taxonomy, a direct comparison between the behavior of Dutch and Paraguayan animals is complicated. The exact species, sex, and age could not be determined from our recordings, whilst individual identification of visiting animals was neither possible in the Netherlands nor in Paraguay. Meijer and Robbers (2014) mostly recorded Old World mice (*Muridae*), whereas most of our mouse visitors belonged to the New World mice (*Cricetidae*) family. Rats were frequently recorded wheel running in the Netherlands, but, although they similarly visited the wheels in Paraguay frequently, there was solely one wheel running occasion of a rat recorded. Even though differences in species diversity cannot be disregarded as one of the most influential factors, the question remains as to what might influence the small mammal (*Rodentia*) communities in different parts of the world to react so differently to an artificial object planted in their environment.

#### Human influence

The difference in probability with which animals encounter humans and their propensity to habituate to human-modified environments may provide a plausible explanation to resolve discrepancy found between wheel running activities at the two locations. The Netherlands and Paraguay differ substantially in population density. Paraguay was extremely sparsely populated with 17 people/km^2^ with respect to 505 people/km^2^ in the Netherlands in 2016 (The World Bank Group, 2017b). Therefore, animals in the Netherlands have a substantially higher chance of encountering people and becoming accustomed to man-made objects than those living in Paraguay. Several rodent species move indoors during unfavorable weather conditions (Frantz and Comings, 1976), thus further increasing the probability of habituation in urban areas. Escaped rodents with previous access to a running wheel will voluntarily run in them when encountered outside captivity (Kavanau, 1967). However, since 1) the nearest village is 3.8 km away from the closest wheel and separated by a permanent lake, and 2) Paraguayan village people do not keep rodents or opossums as pets, it is extremely unlikely that escaped animals, previously exposed to a running wheel, were present in our experimental area.

Urbanization level can directly affect changes in ecological parameters and behavior (Luniak, 2004). Considering that feral animals in urban and semi-urban habitats behave substantially different from wild animals in wild habitats (Fonio et al., 2006; Luniak, 2004), differences in small mammals’ wheel running behavior would be expected. Indeed, when comparing the average number of wheel running cases per month per wheel for the baited phase in the urban (Netherlands), semi-urban (Netherlands), and wild (Paraguay) habitats, a seemingly declining trend in wheel running activity over a gradient from urban to wild habitats emerges (42.1; 12.7 and 0.1 runs per month per wheel, respectively). A similar declining trend emerges when comparing the unbaited phases (4.6; 0.4 runs per month per wheel, respectively for the urban and wild habitats). A comparison with three habitat types in the unbaited phase is not possible as Meijer and Robbers (2014) did not stop providing food in their semi-urban area. Differences in daily activity regimes between Paraguay and the Netherlands further supports the hypothesis that an uneven level of urbanization is potentially responsible for the differences in recorded wheel running activities. As previous research shows, artificial light affects the behavior of terrestrial mice and can alter their circadian rhythm (Bird et al., 2004). At the study sites in Paraguay light pollution is virtually absent and only 1 of 1857 (0.05%) animal visits was observed during daytime. In contrast, Meijer and Robbers (2014) found no significant difference between animal visits during day and night in their urban area, which, as they already suggest, indicates that light pollution affects the rodents’ behavior at their study site. Altogether, since Meijer and Robbers’ (2014) experimental sites are urban and semi-urban, their conclusion that “running in wheels can be a voluntary behavior for *feral* animals in nature” is not necessarily applicable to *wild* animals.

Despite several known hypotheses, e.g., exploration (Mather, 1981), energy regulation (Collier and Leshner, 1967), and feedback dysfunction (Sherwin, 1998), still no consensus regarding the causality of wheel running behavior is reached. On rare occasions, though, wheel running can be pathological in wild environments, a result of abnormal behavior (Mason and Würbel, 2016), toxins (Winrow et al., 2003), or parasites (Hay et al., 1986). Our results demonstrated that small mammals in a remote natural site do not show voluntary wheel running on a similar magnitude that was reported by Meijer and Robbers (2014). However, besides different urbanization levels, taxonomic differences between the mammalian fauna of the two countries and the timing of the studies may still play a role in the observed variation. Since wheel running was observed twice in Paraguay, we cannot state that voluntary wheel running by wild mammals is completely absent in wild environments. However, whether or not the observations in Paraguay show actual wheel running behavior or are merely attempts of animals exploring their surroundings, remains uncertain. The issue of whether wheel running really occurs in the wild, thus, remains unclear and requires additional research.

### Is wheel running a stereotypic behavior?

There are numerous suggested definitions for stereotypic behavior, but all include the same descriptive labels: repetitive, invariant, and devoid of obvious goal or function (Mason, 1991; Mason and Rushen, 2008). When discussing these descriptive definitions, which do not consider the presence or absence of stereotypies in wild environments, wheel running clearly falls into this category (Sherwin, 1998; Richter et al., 2014; Mason and Würbel, 2016). Previous research shows that stereotypic behavior usually indicates that an animal’s psychological welfare is at a suboptimal level (Marriner and Drickamer, 1994). However, whether animals will resort to stereotypic behaviors in the wild when experiencing similar stress-related discomfort is still unclear. Furthermore, if we agree with Meijer and Robbers (2014) who state that “all (authors) agree that stereotypic behavior only occurs in captivity”, and also that wheel running occurs, at least in the Netherlands, in the ‘wild’, then labelling it as a stereotypic behavior would become debatable. Mason and Würbel (2016) argue that “observing wheel running in wild animals does not demonstrate that laboratory animals’ wheel running is normal, because abnormal behaviors often develop from normal ones.” Thus, abnormal behavior can occur in wild animals in wild environments, and this, therefore, further explains the anomaly that stereotypic behaviors can be observed in the wild.

Our results suggest that voluntary wheel running in the wild can occur, at best, with a much lower frequency than Meijer and Robbers (2014) reported from urban and semi-urban habitats. The rarity of recordings, the low velocities, and the short ‘running’ periods may make it even debatable that our recorded cases can be categorized as the same stereotypical wheel running behavior observed in captive rodents or those recorded by Meijer and Robbers (2014). Therefore, since 1) wheel running neatly fits the definition of a stereotypy, 2) it is, on some occasions, possible for stereotypies to occur outside captivity and thus, also in the habitats described by Meijer and Robbers (2014), and 3) no sustained wheel running behavior is observed by truly wild animals in Paraguay, we contend that wheel running is indeed a stereotypic behavior. We suggest that the presence of voluntary wheel running in nature is a result of synurbization and is likely to be absent in truly wild animals whose behavior is not influenced by human activity. This conclusion further strengthens the position of wheel running as being regarded as a stereotypic behavior.

Stereotypies in animals seem to contain similar mechanisms as stereotypies in humans (Mason, 1991; Garner, 2005). By studying the relationship between wild and captive animals’ behavior — more specifically wheel running — we can not only widen our knowledge of animal welfare and enrichment, but also improve our understanding of human psychological and physiological disorders (Mason, 1991; Cotman and Berchtold, 2002; Waters et al., 2013; Richter et al., 2014). With the exception of Meijer and Robbers (2014) and the present study, all previous scientific literature investigating stereotypic behaviors were solely based on captive or wild-caught animals which result in experimental outcomes that are not necessarily applicable to free-living animals (Cooper and Nicol, 1996; Garner, 2005). Further studies must include a focus on wild animals to investigate how habitat type, urban conditions, light pollution, population density, and seasonality influence voluntary wheel running in order to tighten the loose definition of stereotypic behaviors with regards to non-captive animals and to shed light on their underlying natural causes.

## Acknowledgements

We thank Karina Atkinson from Fundacion Para la Tierra for the permission to conduct our research at the Para La Tierra ecological station, Malvina Duarte, the owner of Laguna Blanca, for her foresight and support, Els van Suijlekom for her help conceiving the study, and particularly Bence Schmatovich for the useful comments and discussions about the topic. We also thank the staff members, interns, and volunteers who helped with fieldwork, Eleonóra Fitos for the habitat photographs, and Gabor L. Lovei and Nick A. Littlewood for comments on the manuscript. We would also like to thank the USFW, Lush Cosmetics, and Rolex Awards for Enterprise for their financial support which paid for equipment used during our study.

## Ethical Statement

All work was conducted with the relevant permits necessary for work with wildlife in Paraguay (Secretaria del Ambiente (SEAM); Permiso No 110/2015). No animals were harmed during the experiments.

## Authorship statement

PvL and MAG contributed equally to this work: conceived and designed the study, collected field data, participated in data analysis, and drafted the manuscript; GP participated in the design of the study, provided GIS map, coordinated the study, and helped draft the manuscript; JS contributed to the conception and design, and helped draft the manuscript. All authors gave final approval for publication.

## Supplementary Data

*Supplementary Data S1*. Dataset of all wheel movements categorized chronologically with locations, date, time, taxa, wheel movements, wheel movement durations, and habitats.

*Supplementary Data S2*. Movie clip of wheel running behavior by unidentified opossum (Didelphidae, suspected grey short-tailed opossum *Monodelphis domesitca*) in the transitional forest. Baited period, 24 April 2016, 18:34.

*Supplementary Data S3*. Movie clip of wheel running behavior by unidentified rat in the transitional forest. Unbaited period, 23 May 2016, 18:01. Note that the second smaller wheel present in the video was part of a supplementary project investigating the influence of wheel size on rodent activity. The presence of this supplementary wheel had no statistical influence on the study and the subsequent data were omitted from the present analysis.

*Supplementary Data S4*. Movie clip of wheel movement from inside the wheel (WMI) by unidentified mouse, whilst not running. Preliminary period, 3 March 2016, 00:11.

*Supplementary Data S5*. Movie clip of wheel movement from outside the wheel (WMO) by unidentified rat. Baited period, 18 April 2016, 03:23.

*Supplementary Data S6*. Movie clip of wheel pulling by an unidentified opossum (Didelphidae, suspected *Marmosa sp*.). Preliminary period, 15 March 2016, 01:34.

